# Coupled Dynamics of Behavior and Disease Contagion Among Antagonistic Groups

**DOI:** 10.1101/2020.06.17.157511

**Authors:** Paul E. Smaldino, James Holland Jones

**Affiliations:** University of California, Merced; Stanford University

**Keywords:** transmission dynamics, coupled contagion, homophily, outgroup aversion, social distancing

## Abstract

Disease transmission and behavior change are both fundamentally social phenomena. Behavior change can have profound consequences for disease transmission, and epidemic conditions can favor the more rapid adoption of behavioral innovations. We analyze a simple model of coupled behavior-change and infection in a structured population characterized by homophily and outgroup aversion. Outgroup aversion slows the rate of adoption and can lead to lower rates of adoption in the later-adopting group or even behavioral divergence between groups when outgroup aversion exceeds positive ingroup influence. When disease dynamics are coupled to the behavior-adoption model, a wide variety of outcomes are possible. Homophily can either increase or decrease the final size of the epidemic depending on its relative strength in the two groups and on *R*_0_ for the infection. For example, if the first group is homophilous and the second is not, the second group will have a larger epidemic. Homophily and outgroup aversion can also produce dynamics suggestive of a “second wave” in the first group that follows the peak of the epidemic in the second group. Our simple model reveals dynamics that are suggestive of the processes currently observed under pandemic conditions in culturally and/or politically polarized populations such as the United States.

## 1. Introduction

Behavior can spread through communication and social learning like an infection through a community (Bass, 1969; Centola, 2018). Cavalli-Sforza and Feldman, who pioneered treating cultural transmission in an analogous manner to genetic transmission, noted that “another biological model may offer a more satisfactory interpretation of the diffusion of innovations. The model is that of an epidemic” (Cavalli-Sforza and Feldman, 1981, 32-33). The biological success of *Homo sapiens* has been attributed to its capacity for cumulative culture, and particularly to the rapid and flexible adaptability that arises from social learning (Henrich, 2015). Adoption of adaptive behaviors during an epidemic of an infectious disease could be highly beneficial to both individuals and the population in which they are embedded (Fenichel et al., 2011). Coupling models of behavioral adoption and the transmission of infectious disease, what we call *coupled contagion* models, may thus provide important insights for understanding dynamics and control of epidemics. While we might expect strong selection—both biological and cultural—for adaptive responses to epidemics, complications such as the potentially differing time scales of culture and disease transmission and the existence of social structures that shape adoption may complicate convergence to adaptive behavioral solutions.

In this paper, we explore the joint role of *homophily* —the tendency to form ties with people similar to oneself—and *outgroup aversion*—the tendency to avoid behaviors preferentially associated with an outgroup. Identity exerts a powerful force on the dynamics of behavior (Hogg and Abrams, 2007; Bishop, 2009; Mason, 2018; Smaldino, 2019; Klein, 2020; Moya et al., 2020). This is because identity at least partly determines whom we associate with, communicate with, and strive to either emulate or avoid. Our analysis is predicated on the idea that this matters for the dynamics of infection. For example, Salathé and Bonhoeffer (2008) showed that if rates of vaccine adherence cluster on networks, as when communities collectively adopt identity-based positions on the likely costs and benefits of vaccination (Bauch and Earn, 2004) or when like-minded individuals tend to assort in social networks (Bishop, 2009), the overall vaccination rates needed for herd immunity can be substantially higher than suggested by models that assume random vaccination.

Homophily involves interactions with ingroup members at rates higher than expected by chance. Homophily is often treated as though it were a global propensity for assortment by type (e.g. Centola, 2011). However, homophily is frequently observed to a greater or lesser degree across subgroups, a phenomenon known as differential homophily (Morris, 1991). Consider a case of two interacting groups, where one is more homophilous than the other. The less homophilous group may consist of more “frontline” workers, who are exposed to a broader cross-section of the population by nature of their work. In such cases, differential homophily may lead to differential disease dynamics in each group.

Members of opposed identity groups not only engage with the world differently, they can react in divergent ways to identical stimuli. Asked to watch political debates or hear political arguments, partisans often grow more strongly partisan, to the consternation of moderates (Taber et al., 2009). In the U.S., partisan identities have become increasingly defined in terms of their opposition to the opposing party (Abramowitz and Webster, 2016). When considering the adoption of products, consumers often become disenchanted with otherwise attractive purchases if the products are associated with identity groups viewed as different from their own (Berger and Heath, 2007, 2008). Smaldino et al. (2017) modeled the spread of a behavior among members of two groups who responded positively to the behavioral contagion but tended to reject it if it was overly associated with the outgroup. They showed that outgroup aversion not only decreased the overall rate of adoption, but could also delay or even entirely suppress adoption in one of the groups. While populations vary in the extent to which they are polarized or parochial, identity clearly matters to the adoption of health behaviors in at least communities. For example, in the U.S., people who identify with the right-wing Republican party are much less likely than those identifying with the center-left Democratic party to endorse mask-wearing or belief in its efficacy in preventing disease transmission during the COVID-19 pandemic (van Kessel and Quinn, 2020).

Several previous studies have considered the coupled contagion of behavior and infection, usually focused on cases where the behavior is one that decreases the spread of the disease (such as social distancing or wearing face masks) and sometimes using the assumption that increased disease prevalence promotes the spread of the behavior (Tanaka et al., 2002; Epstein et al., 2008; Funk et al., 2010; Verelst et al., 2016; Fast et al., 2015; Fu et al., 2017; Hébert-Dufresne et al., 2020; Mehta and Rosenberg, 2020). These models typically assume that individuals differ only in behavior and disease status. Thus, the spread of both disease and behavior depend primarily on rates of behavior transmission and disease recovery. This is true even of models in which the population is structured on networks. Network structure can change the dynamics of contagion. However, contrary to the assumptions of most models, behavioral distributions on social networks are anything but random. People assort in highly non-random ways (McPherson et al., 2001) and these non-random associations both drive and are driven by social identity. This suggests that the role of social identity is an important, but under-studied, component of coupled contagion models.

Here, we consider how identity—and particularly homophilous interactions with in-group members and aversion to adopt behaviors used by an outgroup—influences the spread of novel behaviors that consequently affect the transmission of infectious disease. The model we will present is complex, and hence challenging to analyze. To help us make sense of the dynamics, we will first describe the dynamics of infection and behavior adoption in isolation, and then explore the full coupled model. We will first show how homophily can introduce temporal delays in the infection trajectories between groups. We will then show how outgroup aversion can lead to reduced or even fully inhibited behavior adoption by the later-adopting group. Finally, we will analyze the fully coupled model and show how the identity-driven forces we consider can lead differentiated identity groups to experience an epidemic in very different ways.

## 2. The SIR model of infection with homophily

We model infection in a population in which individuals can be in one of three states: Susceptible, Infected, and Recovered. When susceptibles interact with infected individuals, they become infected with a rate equal to the effective transmissibility of the disease, *τ*. Infected individuals recover with a constant probability *ρ* per infection per unit time. This is the well-known SIR model of epidemics (Kermack and McKendrick, 1927; Tolles and Luong, 2020). The baseline model assumes random interactions governed by mass action, and the dynamics are described by well-known differential equations. This model yields the classic dynamics in which the susceptible and recovered populations appear as nearly-mirrored sigmoids, while the rate of infected individuals rises and falls. The threshold for the epidemic is given by the basic reproduction number, *R*_0_, which is a measure of the expected number of secondary cases caused by a single, typical primary case at the outset of an epidemic in a population entirely composed of uninfected individuals. An epidemic occurs when *R*_0_ *>* 1. For the basic SIR model in a closed population, 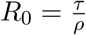.

Our analysis will focus on scenarios where individuals assort based on identity. In this case, assume that individuals all belong to one of two identity groups, indicated with the subscript 1 or 2. Let *w*_*i*_ be the probability that interactions are with one’s ingroup, *i* ∈ {1, 2}. It is therefore a measure of homophily; populations are homophilous when *w*_*i*_ *>* 0.5. It is important to recognize that groups can differ in their homophily (Morris, 1991). For example, if groups differ in socioeconomic class and group 1 tends to employ members of a group 2 as service workers, homophily will be higher for group 1; a member of group 2 is more likely to encounter members of group 1 than the reverse. We can update the equations governing infection dynamics for members of group 1, with analogous equations governing members of group 2.

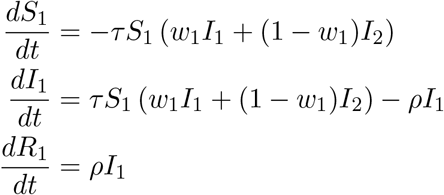

We assume the disease breaks out in one of the two groups, so the initial number of infected in group 1 is small but nonzero, while the initial number of infected in group 2 is exactly zero. Without loss of generality, we have assumed that group 1 is always infected first. When homophily is low, the model exhibits standard SIR dynamics approximating a single unified population. When an infection breaks out in group 1, homophily can delay the outbreak of the epidemic in group 2. Homophily for each group works somewhat synergistically, but the effect is dominated by *w*_2_. This is because the infection spreads rapidly in a homophilous group 1, and if group 2 is not homophilous, its members will rapidly become infected. However, if group 2 is homophilous, its members can avoid the infection for longer, particularly when group 1 is also homophilous. If only group 2 is homophilous, the initial outbreak will be delayed, but the peak infection rate in group 2 can actually be higher than in group 1, as the infection is driven by interactions with both populations (Figure 1).

**Figure 1.**
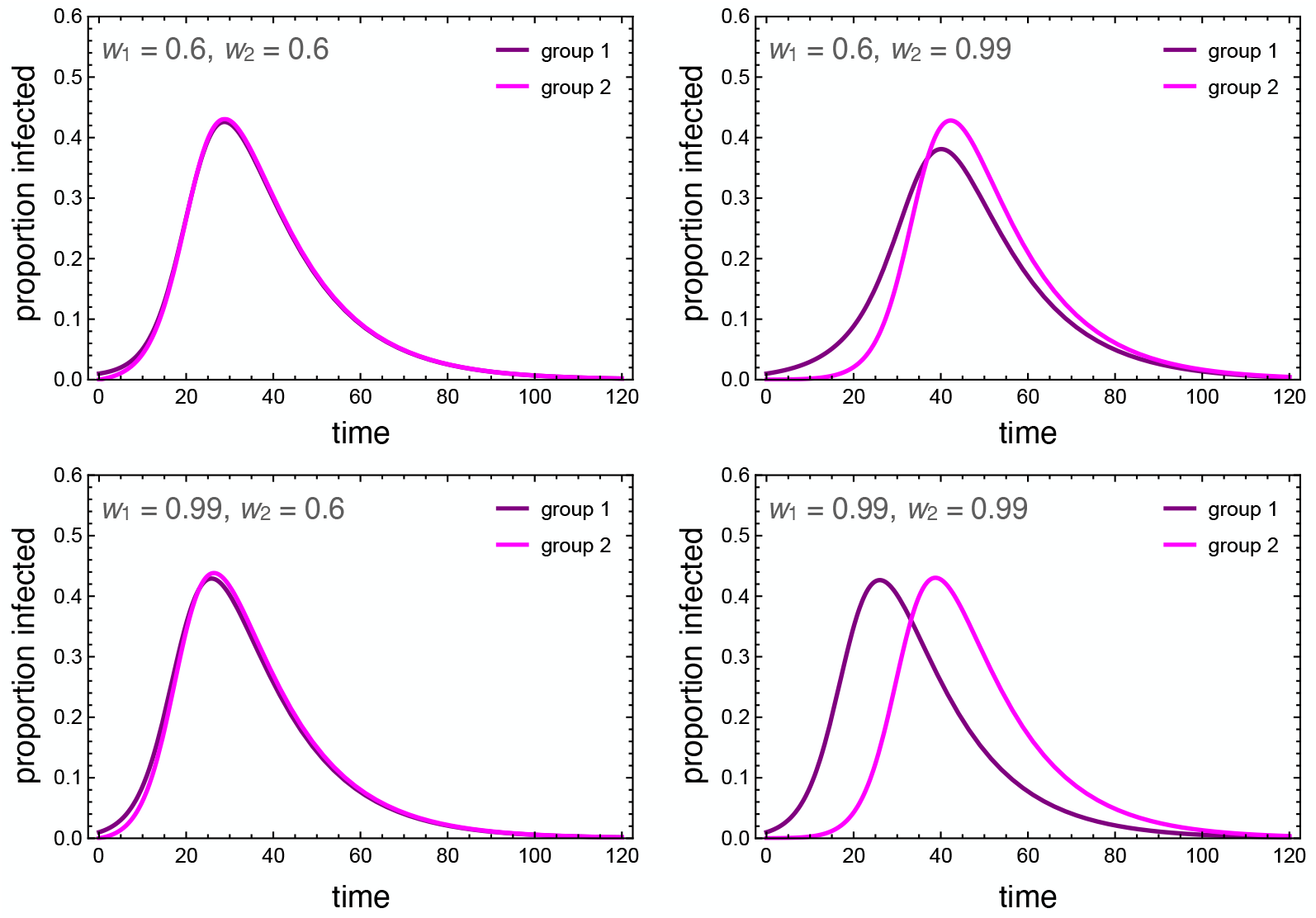
Dynamics of the infected population of each group under low and high homophily (*w*_*i*_ = 0.6, 0.99). Other parameters used were *τ* = 0.3, *ρ* = 0.07, *I*_1_(0) = 0.01, *I*_2_(0) = 0. *R*_0_ ≈ 4.28 in the absence of homophily.

We also considered the case in which the transmissibility of the infection can be reduced to very near the recovery rate, so that *R*_0_ is very close to 1. In this case, homophily can protect groups where infection did not originally break out by keeping members relatively separated from the infection group (Figure S2).

## 3. Behavioral Contagion with Outgroup Aversion

We model behavior adoption as a susceptible-infectious-susceptible (SIS) process, in which individuals can oscillate between adoption and non-adoption of the behavior indefinitely. We view this as more realistic than an SIR process for preventative-but-transient behaviors like social distancing or wearing face masks. To avoid confusion with infection status, we denote individuals who adopted the preventative behavior as Careful (*C*), and those who have not as Uncareful (*U*). Unlike a disease, which is reasonably modeled as equally transmissible between any susceptible-infected pairing, where behavior is concerned, susceptible individuals are more likely to adopt when interacting with in-group adopters, but less likely to adopt when interacting with outgroup adopters. We model the behavioral dynamics for members of group 1 are as follows, with analogous equations^1^ governing members of group 2:

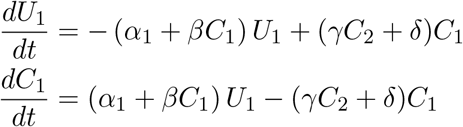

Members of group *i* may spontaneously adopt the behavior independent of direct social influence, and do so at rate *α*_*i*_. This adoption may be due to individual assessment of the behavior’s utility, to influences separate from peer mixing, such as from media sources, or to socioeconomic factors that make behavior adoption more or less easy for certain groups. For these reasons, we assume that groups can differ on their rates of spontaneous adoption. In reality, it is possible for groups to differ on all four model parameters, all of which can influence differences in adoption rates. For simplicity, we restrict our analysis to differences in spontaneous adoption.

Uncareful individuals are positively influenced to become careful by observing careful individuals of their own group, with strength *β*. However, this is countered by the force of outgroup aversion, *γ*, whereby individuals may cease being careful when they observe this behavior among members of the outgroup. The behavior is eventually discarded at rate *δ*, representing financial and/or psychological costs of continuing to adopt preventive behaviors like social distancing or wearing face masks.

This model assumes no explicit homophily in terms of behavioral influence. On the one hand, it seems obvious that we observe and communicate with those in our own group more than other groups. On the other hand, opportunities for observing outgroup behaviors are abundant in a digitally-connected world, which alter the conditions for cultural evolution (Acerbi, 2019). For simplicity, we do not add explicit homophily terms to this system. Instead, we simply adjust the relative strengths of ingroup influence and outgroup aversion, *β/γ*. When this ratio is higher, it reflects stronger homophily for behavioral influence.

Numerical simulations that illustrate the influence of outgroup aversion are depicted in Figure 2. In all cases, the careful behavior is first adopted by group 1. In the absence of outgroup aversion, both groups adopt the behavior at saturation levels, with group 2 being slightly delayed. When outgroup aversion is added, the delay increases, but more importantly, overall adoption declines for both groups. This decline continues as long as the strength of outgroup aversion is less than the strength of positive ingroup influence. A phase transition occurs here (Figure 2C,D). Although group 2 may initially adopt the behavior, adoption is subsequently suppressed, resulting in a polarizing behavior that is abundant in group 1 but nearly absent in group 2.

**Figure 2.**
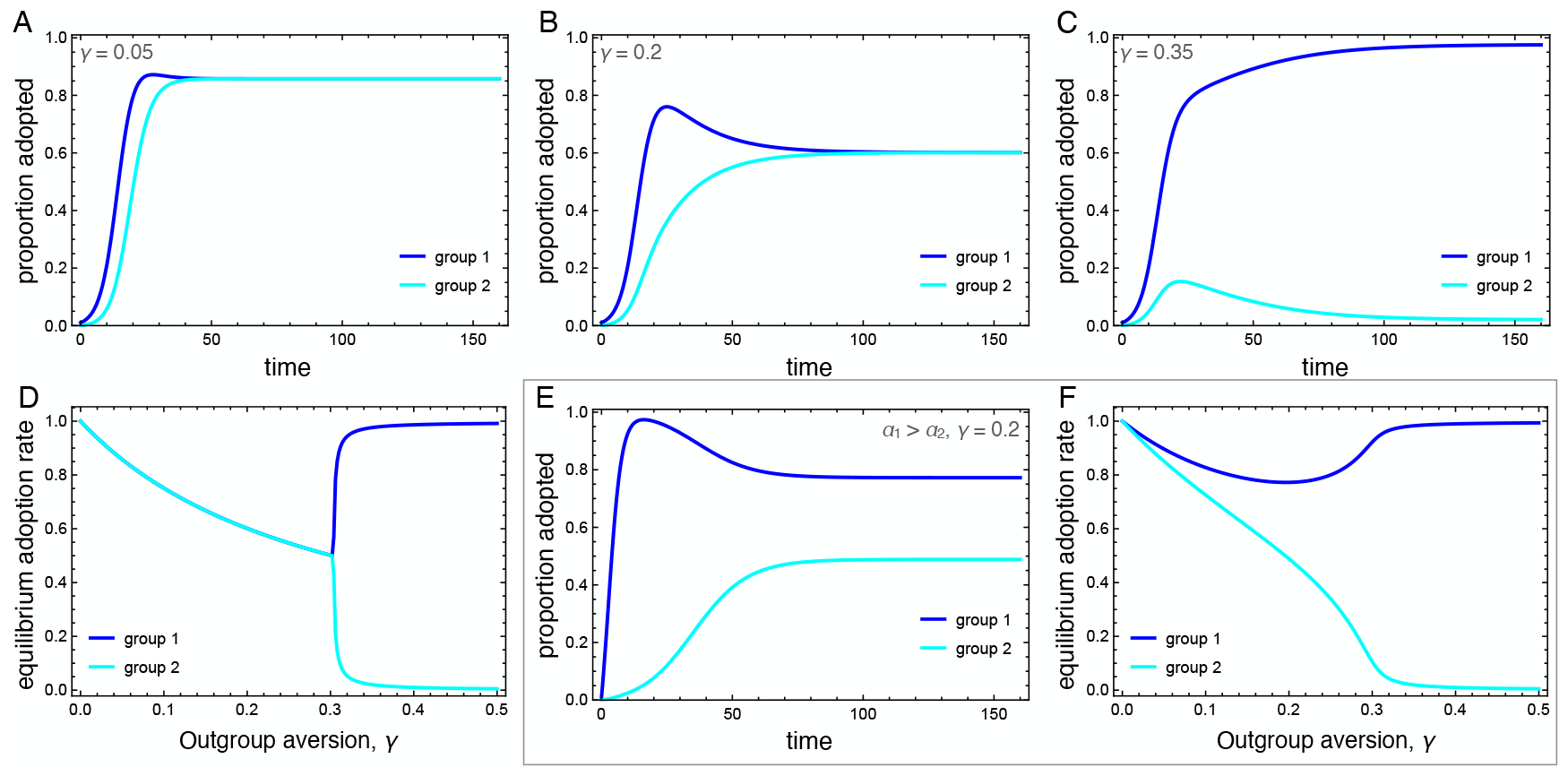
Dynamics of the behavioral adoption. (A-C) Behavior adoption dynamics in each group for different levels of outgroup aversion, *γ*. Parameters used were *α*_1_ = *α*_2_ = 0.001, *β* = 0.3, *δ* = 0, *C*_1_(0) = 0.01, *C*_2_(0) = 0. (D) Equilibrium adoption rates for each group as a function of outgroup aversion, *γ*. A bifurcation occurs when outgroup aversion overpowers the forces of positive influence. (E) Behavior adoption dynamics for *γ* = 0.2 where group 1 has a higher spontaneous adoption rate, *α*_1_ = 0.1. Here, the two groups converge to different equilibrium adoption rates. (F) Equilibrium adoption rates for each group as a function of outgroup aversion, *γ*, when *α*_1_ = 0.1.

We also consider the case in which one group has a higher intrinsic adoption rate, which could be driven by differences in personality types, norms, or media exposure between the two groups. When *α*_1_ *> α*_2_, the equilibrium adoption rate for group 1 could be considerably higher than for group 2, even when ingroup positive influence was greater than outgroup aversion (Figure 2E, F). Note that these differences arise entirely because of outgroup aversion. When *γ* = 0, both groups adopt at maximum levels.

Outgroup aversion has a strong influence on adoption dynamics. It can delay adoption, reduce equilibrium adoption rates, and even suppress adoption entirely in the later-adopting group. As we will see, when the behavior being adopted influences disease transmission, quite interesting dynamics can emerge.

## 4. Coupled Contagion with Homophily and Outgroup Aversion

Before we explore the coupled dynamics of this system, we must add one more consideration to the model. We focus on the adoption of preventative behaviors that decrease the effective transmission rate of the infection, such as social distancing or wearing face masks. We model this by asserting that the transmission rate is *τ*_*C*_ for careful individuals and *τ*_*U*_ for uncareful individuals, such that *τ*_*U*_ ≥ *τ*_*C*_. When considering the interaction between careful and uncareful individuals, we use the geometric mean, so the transmissibility between SU and IU (that is, between susceptible and infected individuals who are both uncareful) is 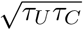. We use the geometric mean so that if either population reduces its transmissibility to zero, transmission among its members becomes impossible.

The full model has six compartments, with two-letter abbreviations denoting the disease and behavioral state (Figure 3). The coupled dynamics for members of group 1 are as follows, with analogous equations governing members of group 2, such that the full system is defined by 12 coupled differential equations. A list of all parameters is presented in Table 1.

**Table 1.**
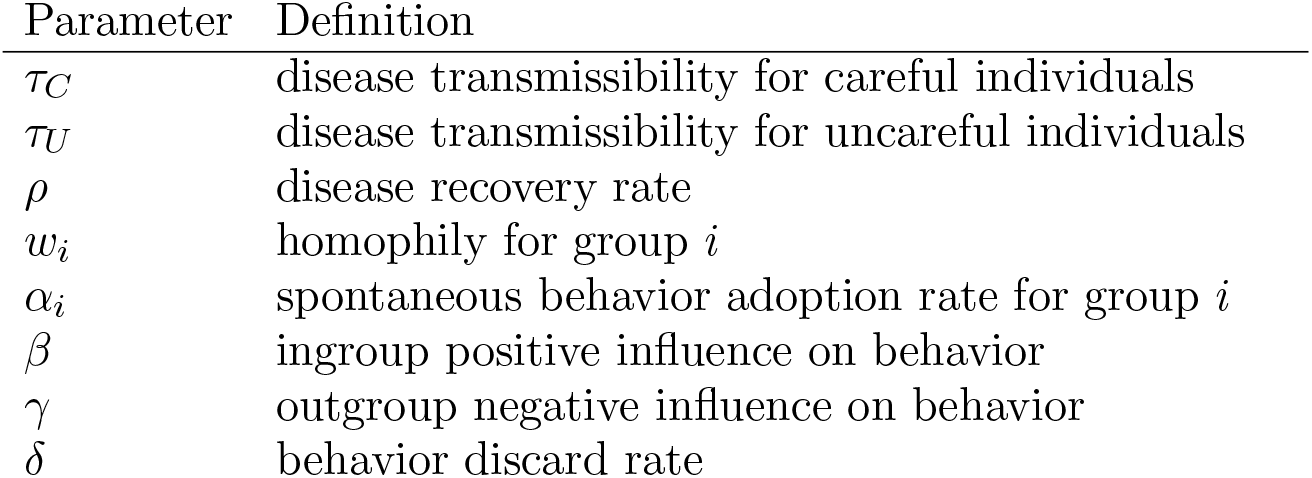
Model parameters.

**Figure 3.**
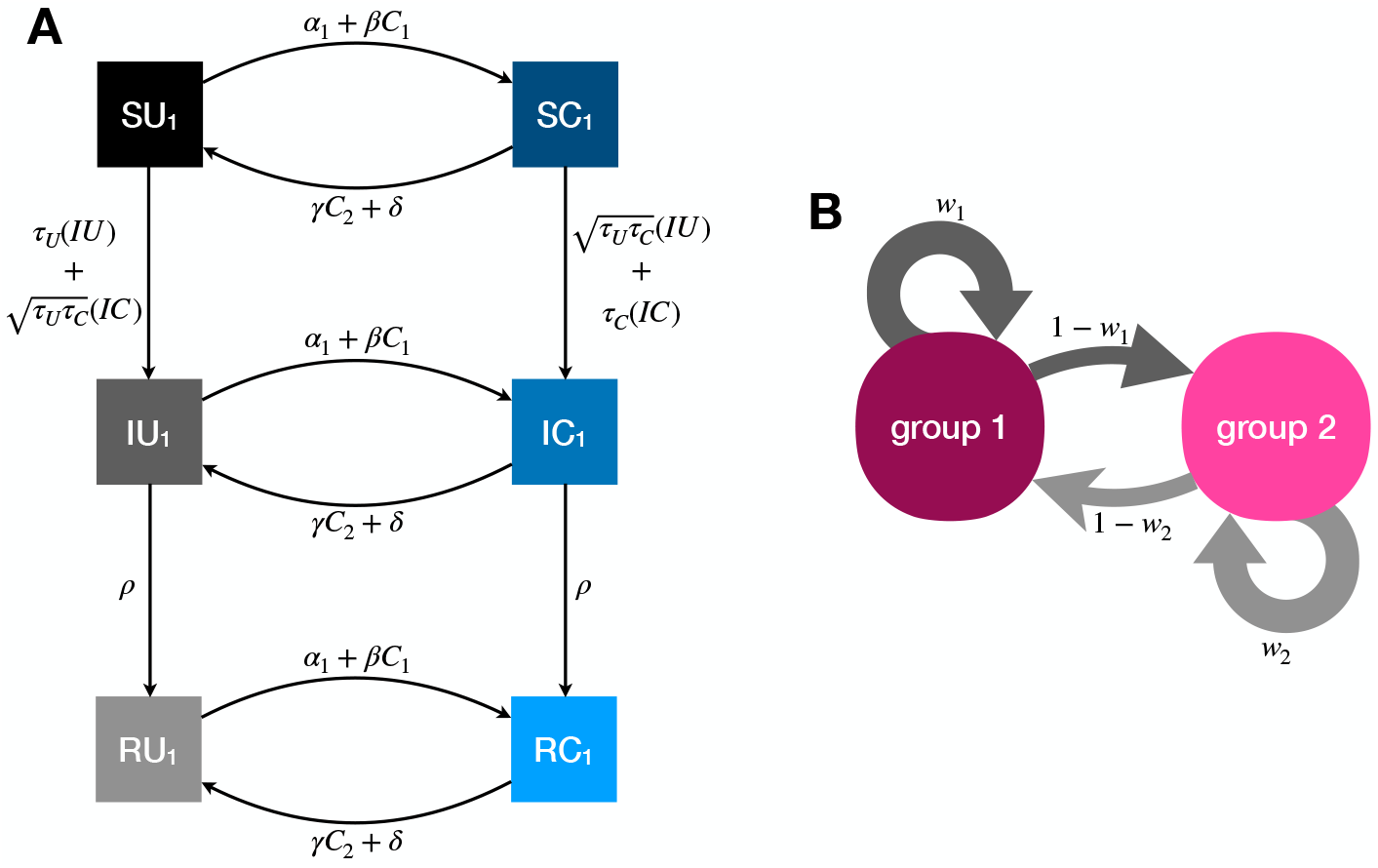
Illustration of the dynamics for the coupled contagion model. (A) Transition probabilities between compartments for members of group 1. For simplicity these probabilities do not include the influence of homophily. (B) homophilous interactions. Members of group *i* have physical contact with members of their own group with probability *w*_*i*_ and members of the outgroup with probability 1 − *w*_*i*_.

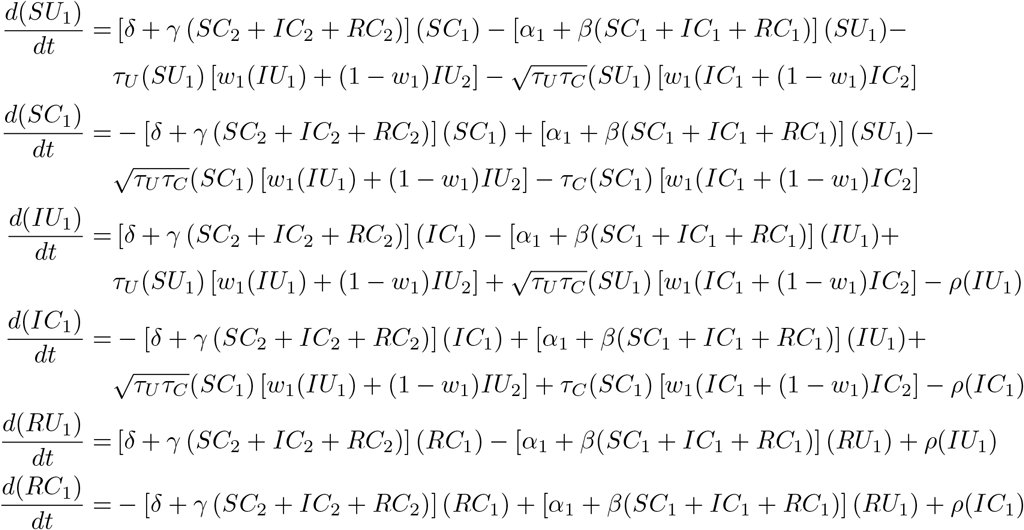

Behavioral adoption is independent of infection status in this model. This may not be a realistic assumptions for some systems, such as Ebola, where the both the infection status of the adopter and the perceived incidence in the population are likely to influence behavior. The assumption seems more realistic for infections like influenza and COVID-19, where infection status is not always transparent and decisions are likely to be made on the basis of more abstract socially-transmitted information. There are intermediate cases, however, such as where media reports of disease prevalence or the perceived availability may influence the adoption of preventative behaviors (Lau et al., 2010; Zhang et al., 2015; Seale et al., 2020). We do not consider such cases here.

To make the behavioral adoption most meaningful, we focus on the case where instantaneous and universal adoption of the careful behavior would decrease the disease transmissibility so that *R*_0_ < 1. That is, if everyone immediately adopted the behavior, the epidemic would fizzle out. However, behavior adoption does not typically work this way. We have already noted that under assumptions of between-group variation and outgroup aversion, a behavior is likely to be adopted neither instantaneously nor universally. The question we tackle now is how those socially-driven facets of behavioral adoption influence disease dynamics.

Figure 4 illustrates the wide range of possible disease dynamics under varying assumptions of homophily and outgroup aversion. A wider range of homophily values are explored in the Supplemental Materials (Figures S4, S5). In the absence of either homophily or outgroup aversion, our results mirror previous work on coupled contagion in which the adoption of inhibitory behaviors reduces peak infection rates, flattening the curve of infection. Due to differences in spontaneous adoption rates, however, group 2 may see a higher peak infection rate even when the infection breaks out in group 1, because the inhibitory behavior spreads more slowly in that group (Figure 4A).

**Figure 4.**
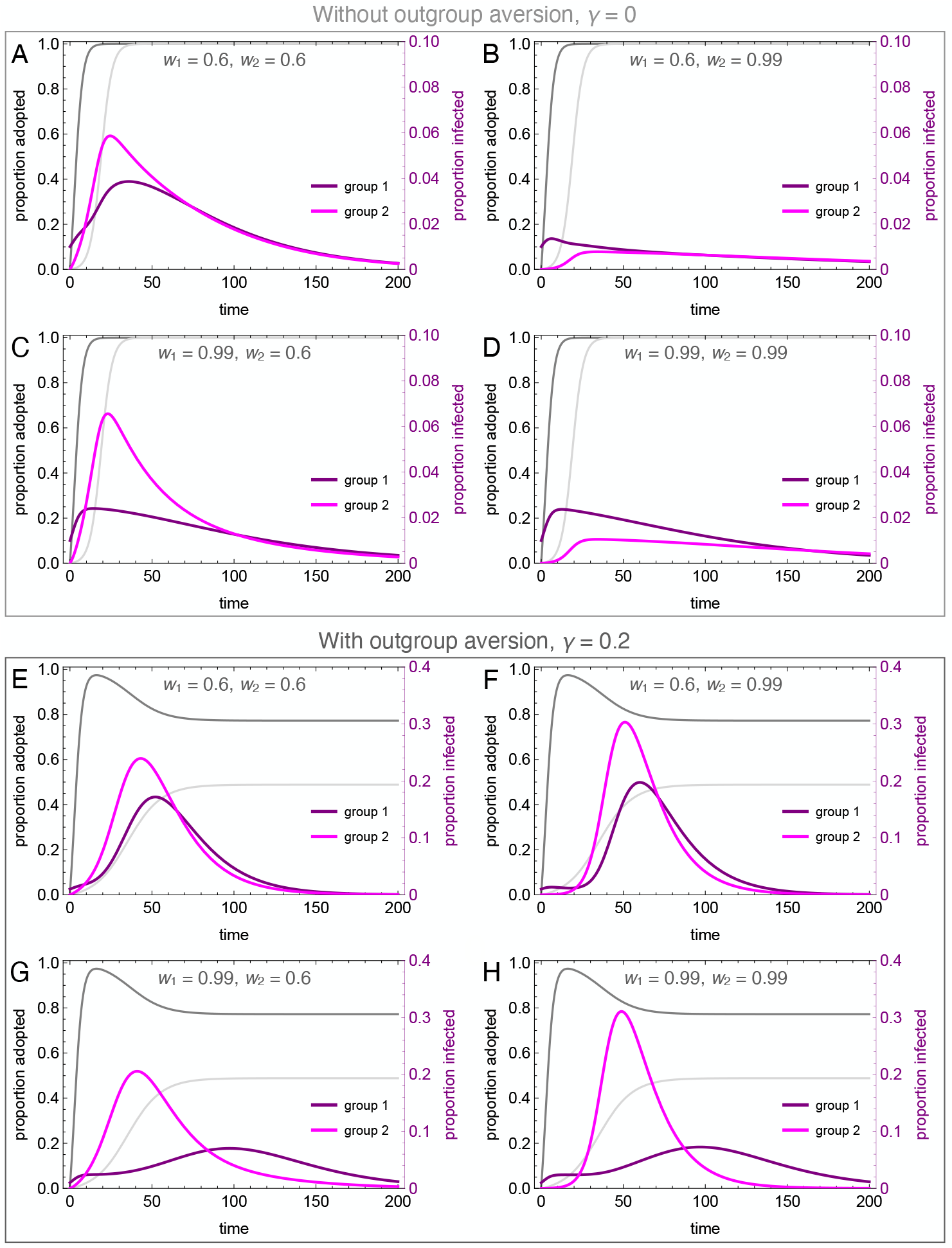
Coupled contagion dynamics when the behavior leads to highly effective reduction in transmissibility, under varying conditions of homophily and outgroup aversion. Notice difference in y-axis scale for infection rate between top and bottom sets of graphs. Parameters used: *τ*_*U*_ = 0.3, *τ*_*C*_ = 0.069, *ρ* = 0.07, *α*_2_ = 0.1, *α*_2_ = 0.001, *β* = 0.3, *δ* = 0, *SU*_1_(0) = 0.98, *SC*_1_(0) = 0.01, *IU*_1_(0) = 0.01, *IC*_1_(0) = *RU*_1_(0) = *RC*_1_(0) = 0, *SU*_2_(0) = 1.0, *SC*_2_(0) = *IU*_2_(0) = *IC*_2_(0) = *RU*_2_(0) = *RC*_2_(0) = 0.

Homophilous interactions further lower infection rates. If group 1 alone is homophilous, the infection rate declines in that group, while peak infections actually increase in group 2 (Figure 4C). This is because group 1 adopts the careful behavior early, decreasing their transmission rate and simultaneously avoiding contact with the less careful members of group 2, who become infected through their frequent contact with group 1. If group 2 alone is homophilous, on the other hand, the infection is staved off even more so than if both groups are homophilous (Figure 4B, D). This is because members of group 2 avoid contact with group 1 until the careful behavior has been widely adopted, while members of group 1 diffuse their interactions with some members of group 2, and these are less likely to lead to new infections.

Outgroup aversion considerably changes these dynamics. First and foremost, outgroup aversion leads to less widespread adoption of careful behaviors, dramatically increasing the size of the epidemic. Moreover, because under many circumstances there will be between-group differences in equilibrium behavior-adoption rates, this can lead to dramatic group differences in infection dynamics. In the absence of outgroup aversion, we saw that homophily in group 2 could lead to an almost total suppression of the epidemic. Not so with outgroup aversion, in which the peak infection rates *increase* relative to the low homophily case (Figure 4E, F). This occurs because homophily causes a delay in the infection onset in group 2. Behavioral adoption slows the epidemic initially in both groups. However, when the infection finally reaches group 1, behavioral adoption has decreased past its maximum due to the outgroup aversion, causing peak infections in both groups to soar.

The dynamics are particularly interesting for the case where the group in which the epidemic first breaks out (group 1 in our analyses) is also strongly homophilous. Due to homophily along with rapid behavior adoption, the epidemic is initially suppressed in this group. However, due to slower and incomplete behavior adoption, the infection spreads rapidly in group 2. As the infection peaks in group 2 while group 1 decreases its behavior adoption rate, we observe a delayed “second wave” of infection in group 1, well after the infection has peaked in group 2 (Figure 4G). This effect is exacerbated when both groups are homophilous, as the epidemic runs rampant in the less careful group 2 (Figure 4H). As shown in the Supplementary Material, the timing of the second wave is also delayed to a greater extent when the adopted behavior is more efficacious at reducing transmission (Figure S6).

We explored the differences in the timing of the infection peaks between the two groups, as illustrated in Figure 5. As noted, homophily in group 1 has a larger effect than homophily in group 2 because the infection first breaks out in group 1. Without outgroup aversion, the infection peak in group 1 is usually closely timed to the infection peak in group 2, usually coming slightly later due to group 2’s lagged adoption of the preventative behavior (Figure 5A). If group 1 has very strong homophily, however, the infection can peak earlier there, as its spread to group 2 is impeded. When outgroup aversion is strong, however, group 2’s adoption of the preventative behavior is severely impeded, which cases its infection rate to peak much earlier than in group 1, and this effect is only bolstered by strong homophily in group 1 (Figure 5B). The effect of outgroup aversion on the differential timing between groups of infection rate peaks is non-monotonic (Figure 5C), peaking at intermediate values of *γ*.

**Figure 5.**
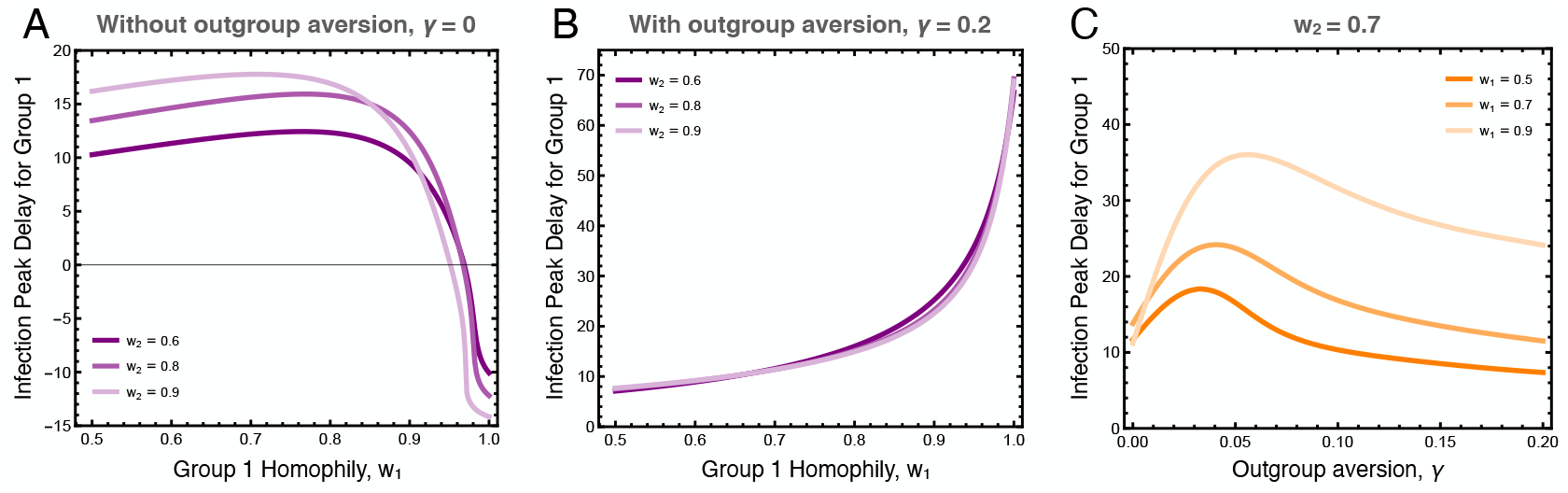
Difference in the timing of the peak infection rates between groups. These plots show the extend to which the peak in group 1 lags behind the peak in group 2. The first two plots show the peak delay for group 1 as a function of group 1 homophily, (A) with and (B) without outgroup aversion, *γ*. The third plot (C) more systematically varies outgroup aversion, for several values of group 1 homophily and moderate group 2 homophily, *w*_2_ = 0.7. Other parameters used: *τ*_*U*_ = 0.3, *τ*_*C*_ = 0.069, *ρ* = 0.07, *α*_2_ = 0.1, *α*_2_ = 0.001, *β* = 0.3, *δ* = 0.

## 5. Discussion

It is well known that disease transmission is influenced by behavior. What is often overlooked is how behavior itself changes within heterogeneous cultural populations. Both population structure and social identity influence who interacts with whom, affecting disease transmission, and who learns from whom, affecting behavior change. We have highlighted two of these forces—homophily and outgroup aversion—and shown their dramatic influence on disease dynamics in a simple model.

In terms of social interaction and behavior adoption dynamics, group identity exerts its influence by way of homophily, a powerful social force. Aral et al. (2009), for example, showed that homophily accounted for more than 50% of contagion in a natural experiment on behavioral adoption. The effect of homophily on diffusion dynamics can be variable. For example, homophily can slow down convergence toward best responses in strategic networks (Golub and Jackson, 2012). This can be critical when the time scales of learning and infection are different. Homophily can also lower the threshold for desirability (or the selective advantage) required for adoption of a behavior. Creanza and Feldman (2014) showed that homophily and selection can have balancing effects—the selective advantage of a trait does not need to be as high to spread when it is transmitted assortatively by its bearers. In the case of our coupled-contagion model, strong homophily interferes with the adaptive adoption of protective behavior. Centola (2011) showed that homophily can increase the rate of adoption of health behaviors, but his experimental population could assort only on positive cues, and had no ability to signal or perceive group identity.

Consider the observed adoption dynamics under differential homophily. When the homophily of group 1 is less than group 2, group 1 can be interpreted as “frontline” workers, who are exposed to a broader cross-section of the population by nature of their work. Outgroup avoidance of this group’s adopted protective behavior can arise if there are status differentials across the groups. Prestige bias, the tendency to adopt behaviors associated with high-status individuals, is a mechanism that can drive differential uptake of novel behavior by different groups (Boyd and Richerson, 1985), for which there is quite broad support (Jiménez and Mesoudi, 2019). When both groups are highly homophilous and outgroup aversion is strong, the resulting dynamics suggest the case of negative partisanship, a type of outgroup aversion in which partisans select actions based not on explicit policy preferences but in opposition to the outgroup (Abramowitz and Webster, 2016). In this case, differences in the relative size of the epidemic will be driven purely by differences in the rates of preventative behavior adoption by the two groups, including those differences induced by outgroup aversion.

Incorporating adaptive behavior into epidemic models has been shown to significantly alter dynamics (Fenichel et al., 2011). Prevalence-elastic behaviors (Funk et al., 2010) are those behaviors that increase with the growth of an epidemic. While these behaviors may be protective, they can also lead to cycling of incidence, which can prolong epidemics. Similarly, the adoption of some putatively-protective behaviors that are actually ineffective can be driven by the existence of an epidemic when the cost of adoption is sufficiently low (Tanaka et al., 2009). We have shown in this paper that group-identity processes can have large effects, leading groups that would otherwise respond adaptively to the threat of an epidemic to behave in ways that put them, and the broader populations in which they are embedded, at risk.

The context of the ongoing COVID-19 pandemic provides some interesting and timely perspective on the relationship between behavior, adaptive or otherwise, and transmission dynamics. While there remains much uncertainty about the infection fatality ratio of COVID-19, and how this varies according to individual, social, and environmental context, it is clear that the great majority of infections do not lead to death (Russell et al., 2020; Meyerowitz-Katz and Merone, 2020). Furthermore, the extensive presymptomatic (or even asymptomatic) transmission of the SARS-CoV-2 (He et al., 2020; Li et al., 2020; Arons et al., 2020) is likely to reduce associations between behavior and local infection rates. We expect that such a situation will not induce strong prevalence-elastic behavioral responses, and that the sorts of identity-based responses we describe here will dominate the behavioral effects on transmission.

How do we intervene in a way to offset the pernicious effects of negative partisanship on the adoption of adaptive behavior? While it may seem obvious, strategies for spreading efficacious protective behaviors in a highly-structured population with strong outgroup aversion will require weakening the association between protective behaviors and particular subgroups of the population. Given that we are writing this during a global pandemic in which perceptions and behaviors are highly polarized along partisan lines, attempts to mitigate partisanship in adaptive behavioral responses seem paramount to support.

The models we have analyzed in this paper are broad simplifications of the coupled dynamics of behavior-change and infection. It would therefore be imprudent to use them to make specific predictions. The goal of this approach is to develop strategic models in the sense of Holling (1966), sacrificing precision and some realism for general understanding of the potential interactions between social structure, outgroup aversion, and coupled contagion (Levins, 1966; Smaldino, 2017). Such models provide a scaffold for the development of richer theories concerning coupled disease and behavioral contagions.

## Acknowledgments

We thank two anonymous reviewers for helpful comments.

## Author Contributions

PES led the work on model conceptualization and analysis, with extensive contribution from JHJ. Both authors wrote and edited the manuscript.

## Research Transparency and Reproducibility

n/a

## APPENDIX (ONLINE SUPPLEMENT)

### Appendix A. The SIR model with homophily

We extended the SIR model to explore scenarios where individuals assort based on identity, as described in the main text. Here we present some additional analyses of this model.

Figure S1 illustrates that when an infection breaks out in group 1, homophily can delay the outbreak of the epidemic in group 2. Homophily for each group works somewhat synergistically, but the effect is dominated by *w*_2_. This is because the infection spreads rapidly in a homophilous group 1, and if group 2 is not homophilous its members will rapidly become infected. However, if group 2 is homophilous, its members can avoid the infection for longer, particularly when group 1 is also homophilous.

**Figure S1.**
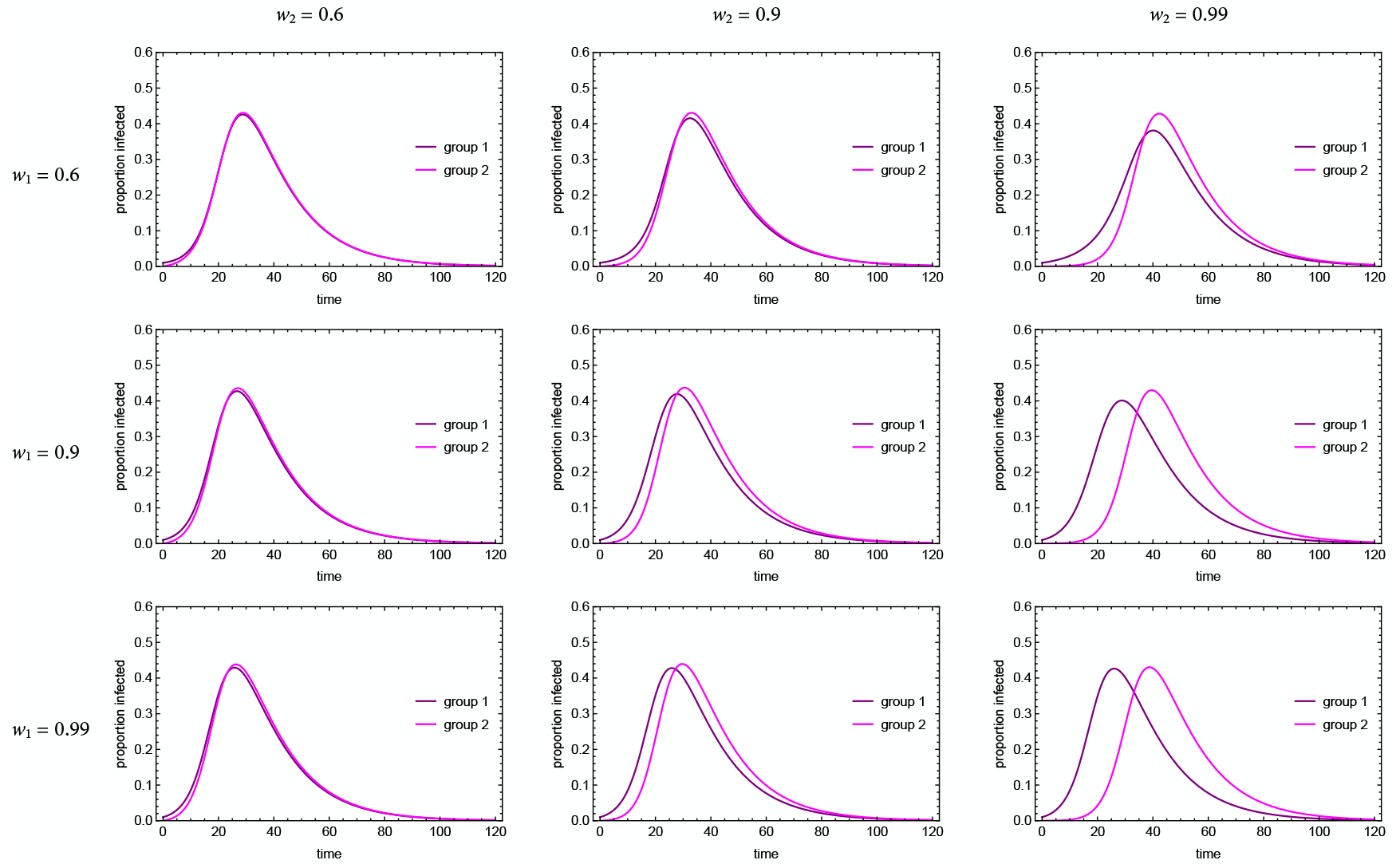
Infection dynamics in the SIR model with asymmetric homophily. Here *τ* = 0.3, *ρ* = 0.07.

We also explored a scenario where *R*_0_ for the basic model was very close to 1, indicating a small epidemic (we used *R*_0_ = 1.14; Figure S2). Note that this calculation of *R*_0_ does not account for homophily; we derive *R*_0_ for the homophily model in the SI Appendix and show that this is a reasonable approximation. When homophily was low (*w* = 0.6), the populations mixed a lot. The proportion of infected individuals in group 1 briefly fell, as the majority of new infected individuals were in group 2. However, the groups quickly matched their pace and experienced the outbreak in tandem. When homophily was high (*w* = 0.99), not only did group 2 experience a delayed outbreak, it also experienced a substantially lower peak infection rate, because the total number of infected individuals at the start of its outbreak was so much lower than that experienced by group 1. Thus, homophily can serve not only to delay an epidemic, but also to reduce it in the cases of lower transmissibility infections.

**Figure S2.**
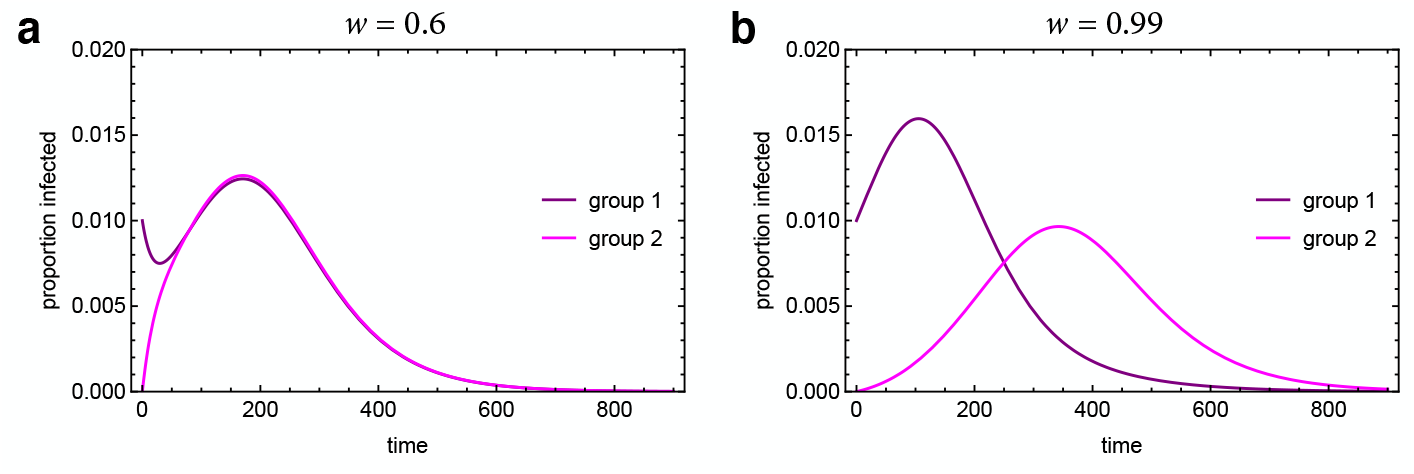
Infection dynamics in the SIR model with homophily when *R*) is close to 1. Here *τ* = 0.08, *ρ* = 0.07, *w*_1_ = *w*_2_ = *w*.

### Appendix B. Basic Reproduction Number

We can calculate the basic reproduction number, *R*_0_, for the homophily model. We employ the next-generation matrix approach described by Heffernan et al. (2005), which concisely summarizes the ideas for calculating *R*_0_ in structured populations articulated by, e.g., Diekmann et al. (1990) and van den Driessche and Watmough (2002).

Following the notation of Heffernan et al. (2005), the next generation matrix **G** is comprised of two component matrices: ***F*** and ***V*** ^−1^, where

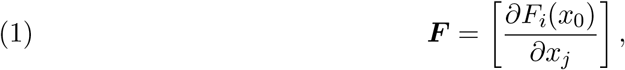

and

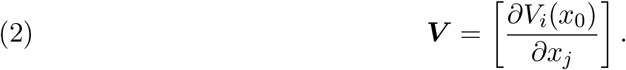

These are square matrices of the partial derivatives of new infections (*F*_*i*_) and transfers between different compartments (*V*_*i*_). The rank of these matrices is the number of distinct classes of infections. *x*_0_ is the disease-free equilibrium state. This matrix should be non-negative, irreducible, and primitive.

is given by the dominant eigenvalue of the matrix ***G*** = ***F V*** ^−1^.

For the homophily model, the only two equations that yield new infections are those for *İ*_1_ and *İ*_2_:

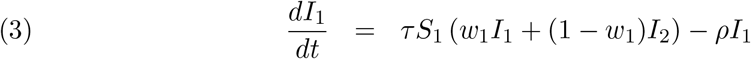

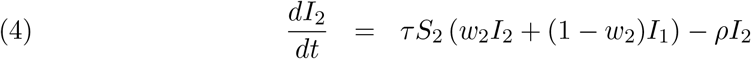

Applying the next-generation-matrix approach described above to these equations, and noting that in the disease-free equilibrium *S*_1_ + *S*_2_ = 1, we get the next-generation matrix:

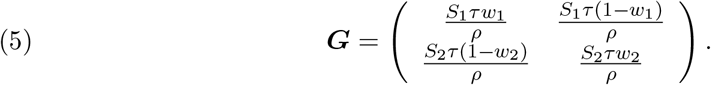

Letting *S*_2_ = 1 − *S*_1_ in the disease-free equilibrium, the larger of the two eigenvalues of this matrix is:

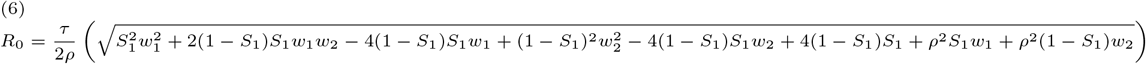

This relationship is greatly simplified by assuming uniform homophily (*w*_1_ = *w*_2_ = *w*):

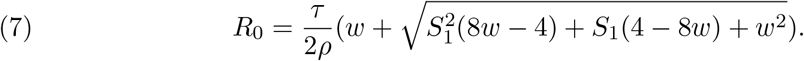

Note that if we collapse the structure of the population such that *S*_1_ = 1 (which also implies that *w* = 1), then equation 7 reduces to *R*_0_ = *τ/ρ*, the standard definition for the basic reproduction number in an unstructured SIR model (Keeling and Rohani, 2007).

We see from figure S3 that structure and homophily (in the absence of coupled adaptive behavior and outgroup aversion) are actually somewhat protective from an epidemic perspective. *R*_0_ is lowest when the population is evenly split between the two groups and when homophily is extreme. This makes sense since structure generally slows epidemics by subdividing the potential for contacts and thereby slowing mixing (Arthur et al., 2017).

### Appendix C. Coupled contagion dynamics

Here we present an extended version of the full model analysis presented in the main text, that includes intermediate homophily of *w*_*i*_ = 0.9. Analysis with no outgroup aversion is shown in Figure S4, and with outgroup aversion is shown in Figure S5. The figures illustrate how homophily and outgroup aversion can interact to produce unintuitive dynamics. When both forces are present, an infection that begins in group 1 can peak earlier and stronger in group 2, followed by a smaller peak in the group where it began.

**Figure S3.**
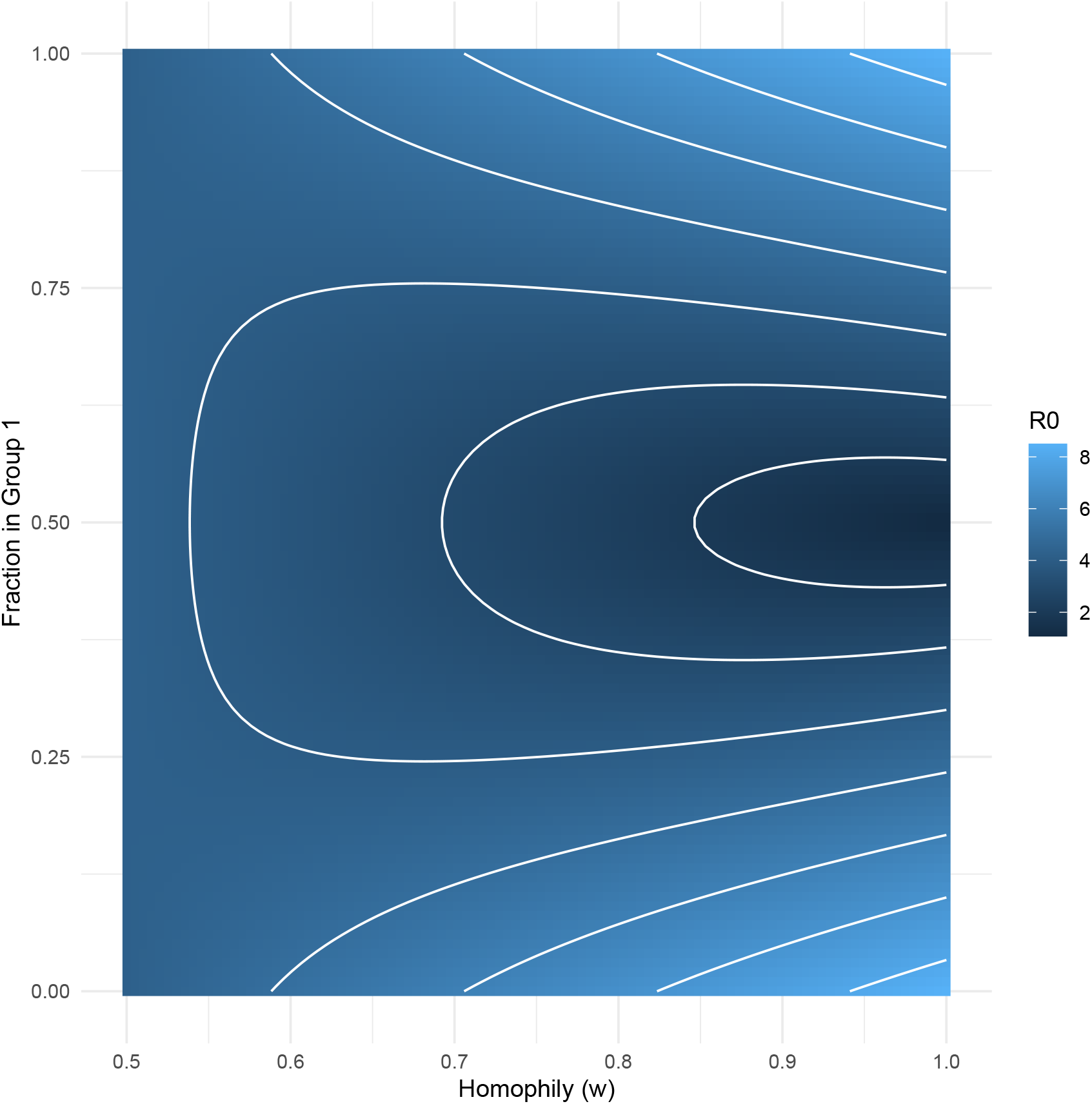
*R*_0_ for the uniform-homophily model (Equation 7) as a function of the strength of homophily (*w*) and the initial population structure. *τ* = 0.3, *ρ* = 0.07.

### Appendix D. Analysis of behavioral efficacy

In the main text analysis, we assumed that the adopted behavior reduced the transmission to below the threshold for *R*_0_ < 1. In other words, if everyone immediately and universally adopted the behavior at the start of the outbreak, it would not become an epidemic. Although we view this as a reasonable assumption (that is, the efficacy of the behavior is reasonable, not the expectation that it will be either immediately or universally adopted), it is also worth examining what happens with the spread of behaviors that reduce transmission, but not below epidemic levels. Figure S6 illustrates the model dynamics for varying levels of behavior efficacy (*τ*_*C*_) with and without outgroup aversion and for both weak and strong homophily.

**Figure S4.**
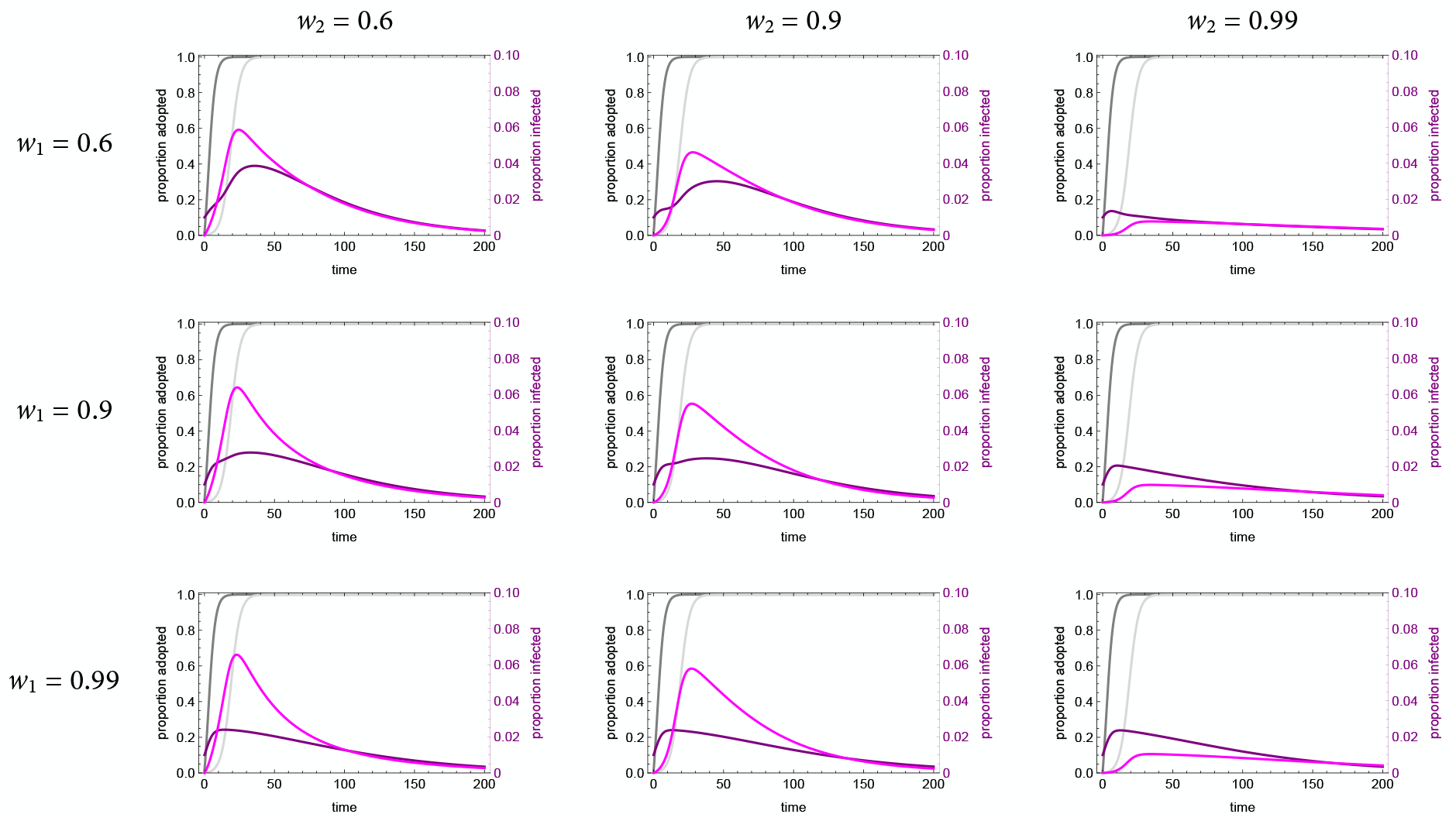
Coupled dynamics of the full model without outgroup aversion (*γ* = 0) for with varying homophily. Darker lines are group 1, lighter lines are group 2. Parameters used: *τ*_*U*_ = 0.3, *τ*_*C*_ = 0.069, *ρ* = 0.07, *α*_2_ = 0.1, *α*_2_ = 0.001, *β* = 0.3, *δ* = 0.

Without outgroup aversion (*γ* = 0), the effect is clear: the more efficacious the behavior, the smaller the epidemic. This occurs because the behavior spreads effectively. With outgroup aversion, two things happen. First, the more effectively the behavior reduces transmission (that is, the smaller *τ*_*C*_ is), the smaller the overall epidemic, but with an effect that is much stronger in group 1. In group 2, the effect of increased behavior efficacy is relatively small, because adoption is reduced and delayed. Second, the better the behavior reduces transmission, the bigger the delay in when group 1 experiences a “second wave.” This illustrates that the dynamics of disease transmission can become quite complex when even simple assumptions about behavior and group structure are considered.

**Figure S5.**
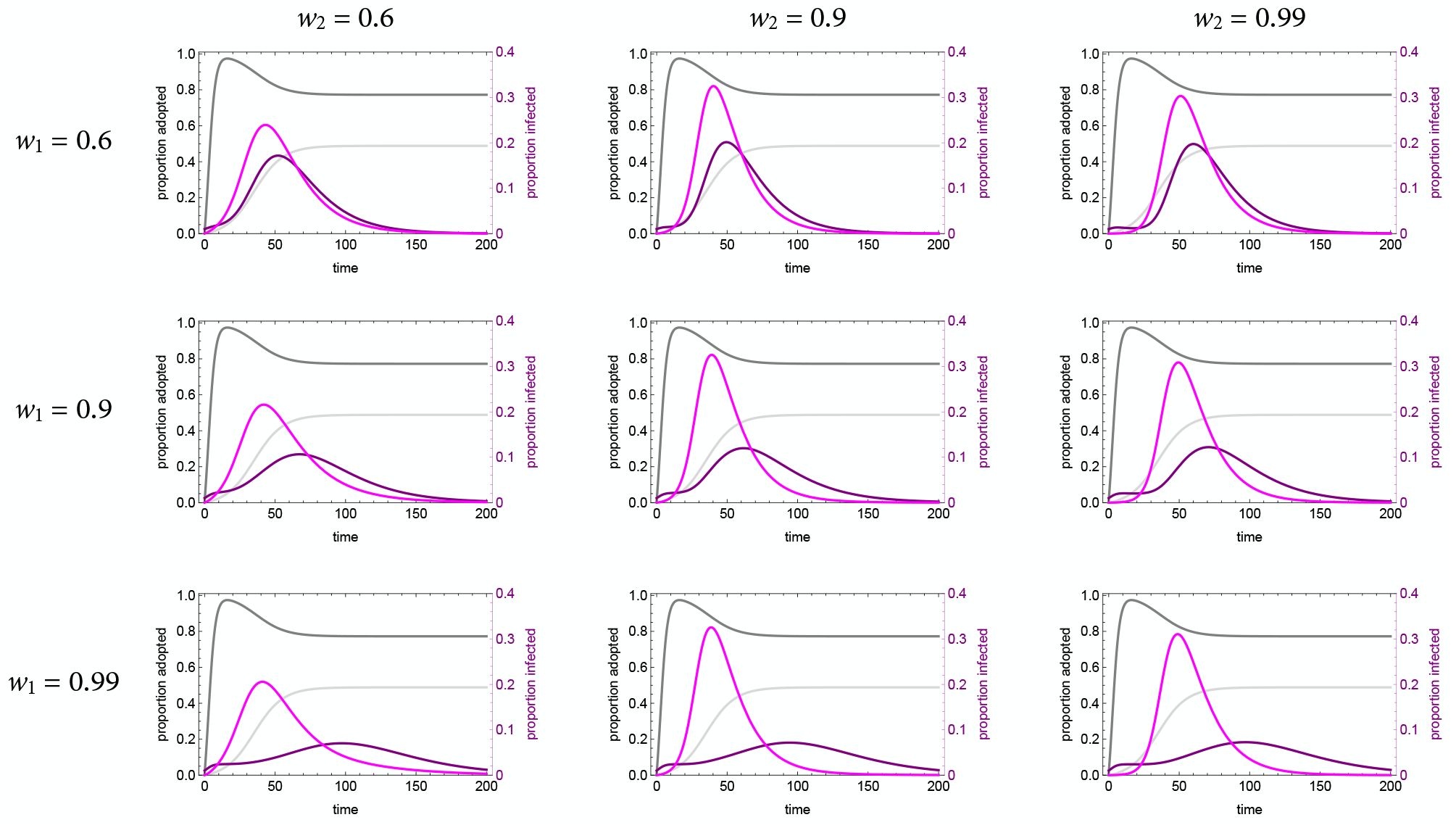
Coupled dynamics of the full model with outgroup aversion (*γ* = 0.2) for with varying homophily. Darker lines are group 1, lighter lines are group 2. Parameters used: *τ*_*U*_ = 0.3, *τ*_*C*_ = 0.069, *ρ* = 0.07, *α*_2_ = 0.1, *α*_2_ = 0.001, *β* = 0.3, *δ* = 0.

**Figure S6.**
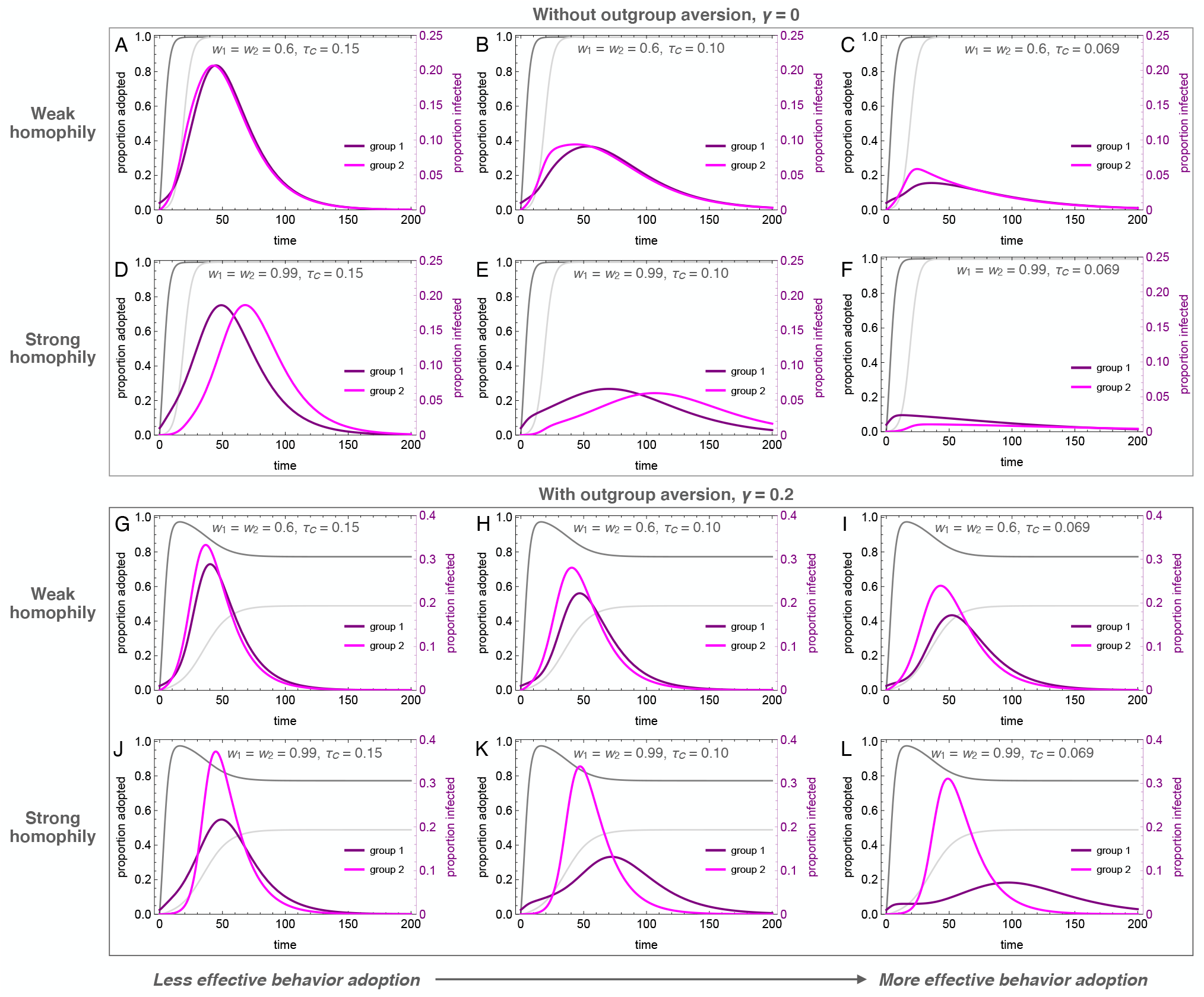
Coupled dynamics of the full model for varying levels of behavior efficacy, *τ*_*C*_ = {0.15, 0.1, 0.069}, where only the last case would provide *R*_0_ < 1 if immediately and universally adopted at the start of the outbreak. We provide analyses with and without outgroup aversion and for both weak and strong homophily. Darker lines are group 1, lighter lines are group 2. Parameters used: *τ*_*U*_ = 0.3, *ρ* = 0.07, *α*_2_ = 0.1, *α*_2_ = 0.001, *β* = 0.3, *δ* = 0.

## Footnotes

1 Because all individuals have either adopted or not, *U*_1_ = 1 − *C*_1_, these coupled equations can be replaced by a single equation through substitutions. For intuitive reasons, we leave them as two coupled equations.

